## Supplementary Information for "Lowland plant arrival in alpine ecosystems facilitates a decrease in soil carbon content under experimental climate warming"

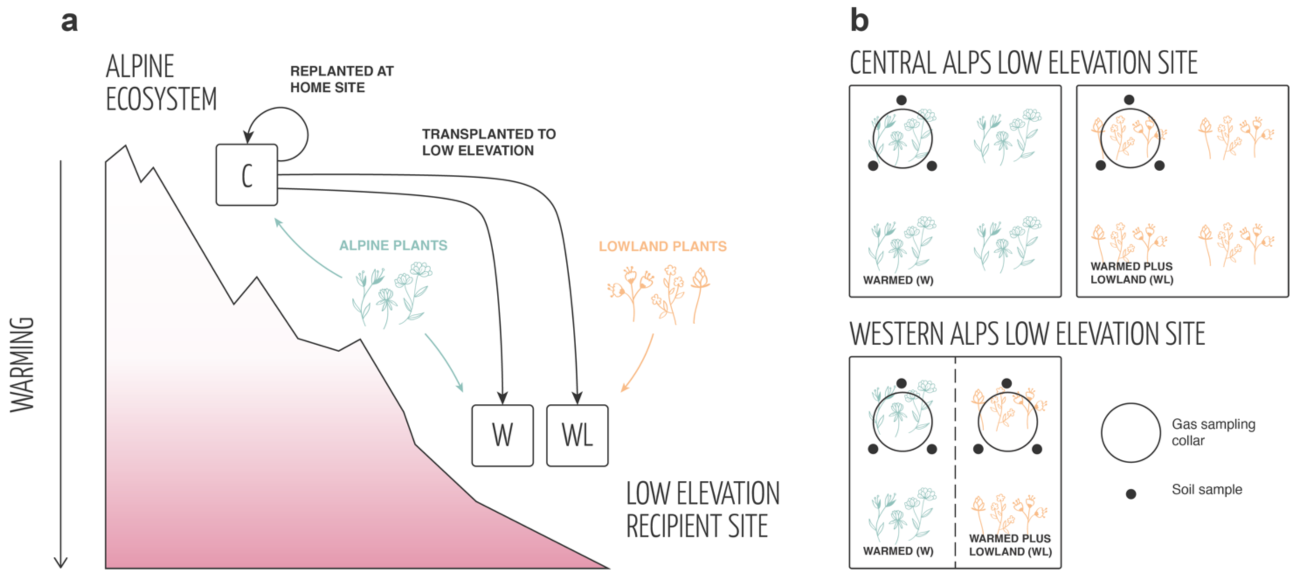

**Fig. S1 | Experimental design.** (a) In both field experiments, intact alpine turfs containing alpine plant communities plus rhizosphere soil were removed from their high elevation home sites and either replanted at the home site (negative control; C) or transplanted to a low elevation recipient site to simulate climate warming (W, WL). At low elevation, half of the turfs were planted with a low abundance of lowland plant species to additionally simulate the arrival of migrating plant species in the ecosystem (WL). Both other treatments (i.e. C, W) were planted with the same abundance of alpine plant species as a disturbance control. (b) The layout of W and WL treatments at the low elevation site differed between western and central alps experiments. In the central alps, W and WL treatments were established in distinct turfs planted adjacently in a block design (N = 10). In the western alps, W and WL treatments were established in different halves of the same turf in a split-plot design (N = 10).

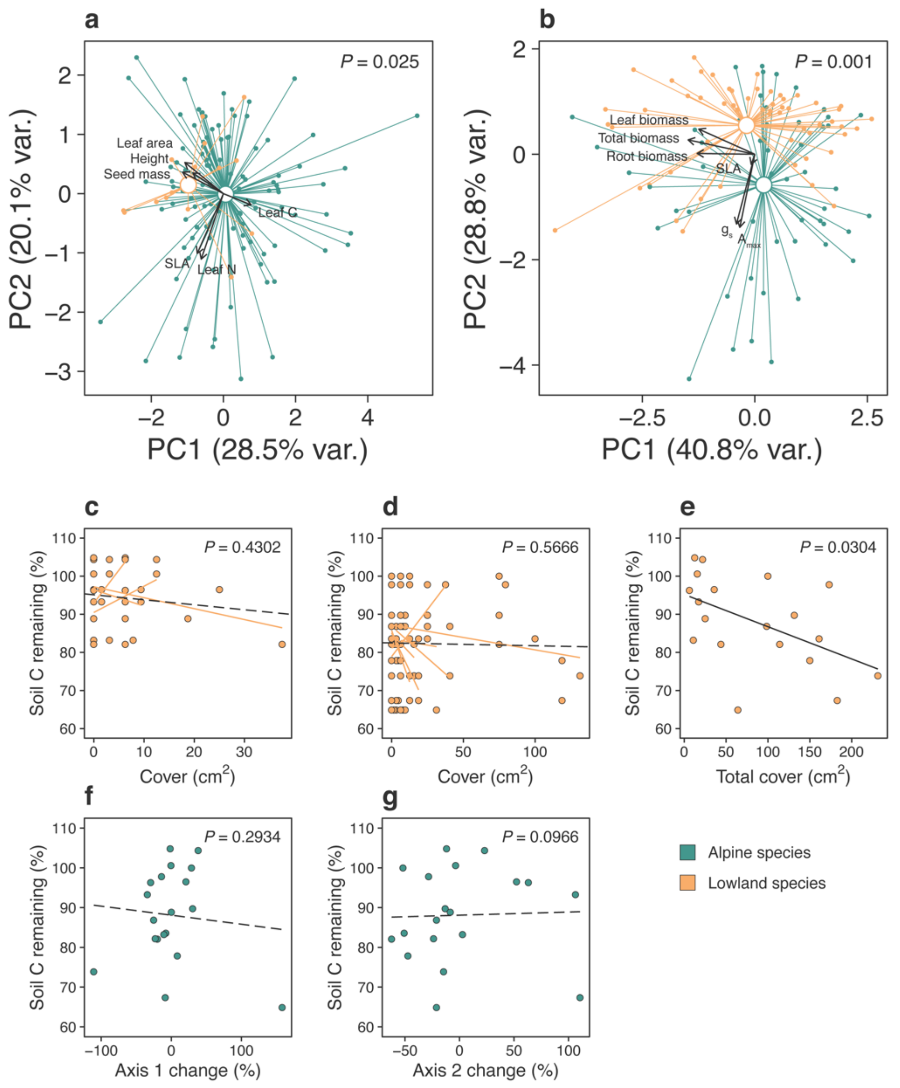

**Fig. S2 | Alpine and lowland plant effects on soil carbon loss.** (a,b) Trait separation of lowland (yellow) and alpine (green) plant species, displayed as PC1 and PC2 scores from PCAs containing (a) TRY data for species in field experiments (N = 242; $F_{1,240}$ = 3.98, *P* = 0.003) or (b) measured data from a glasshouse experiment on selected species (N = 109; $F_{1,107}$ = 17.69, *P* = 0.001). Arrows show loadings and *P*-values refer to alpine-lowland comparison (PERMANOVAs). (c-g) Relationships between soil carbon loss (% remaining) and cover (cm2) of each lowland plant species in the (c) western Alps (N = 36; LME: LR = 0.62, *P* = 0.4302) and (d) central Alps (N = 80; LME: LR = 0.33, *P* = 0.5666) experiments, (e) total lowland plant cover ($cm^2$; N = 19; see Main Text) and (f,g) alpine plant community composition (% change in NMDS axes; N = 18; linear models: NMDS #1: $F_{1,14}$ = 1.19, *P* = 0.2934, NMDS #2: $F_{1,14}$ = 3.17, *P* = 0.0966; region: $F_{1,14}$ = 1.29, *P* = 0.2755). For (c,d), non-significant species-wise fit lines (yellow) are also displayed.

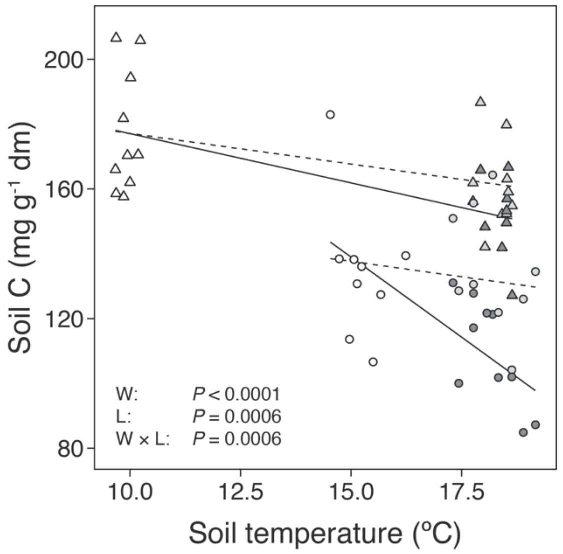

**Fig. S3 | Soil temperature and lowland plant effects on alpine soil carbon loss.** Relationship between soil temperature (ºC) and soil carbon content (C; mg $g^{-1}$ dry mass) in alpine turfs exposed to warming (W; light grey), warming plus lowland plants (WL; dark grey) or an ambient control (C; white). Statistics describe effects of warming (W; LR = 17.65, $P < 0.0001$), lowland plants (L; LR = 11.77, $P = 0.0006$) and their interaction (W × L; LR = 11.72, $P = 0.0006$) from one linear mixed effects model (N = 78), but are visualized with separate fit lines (including slopes) for C-W (dashed lines) and C-WL (solid lines) comparisons in the western Alps (triangles) and central Alps (circles) field experiments.

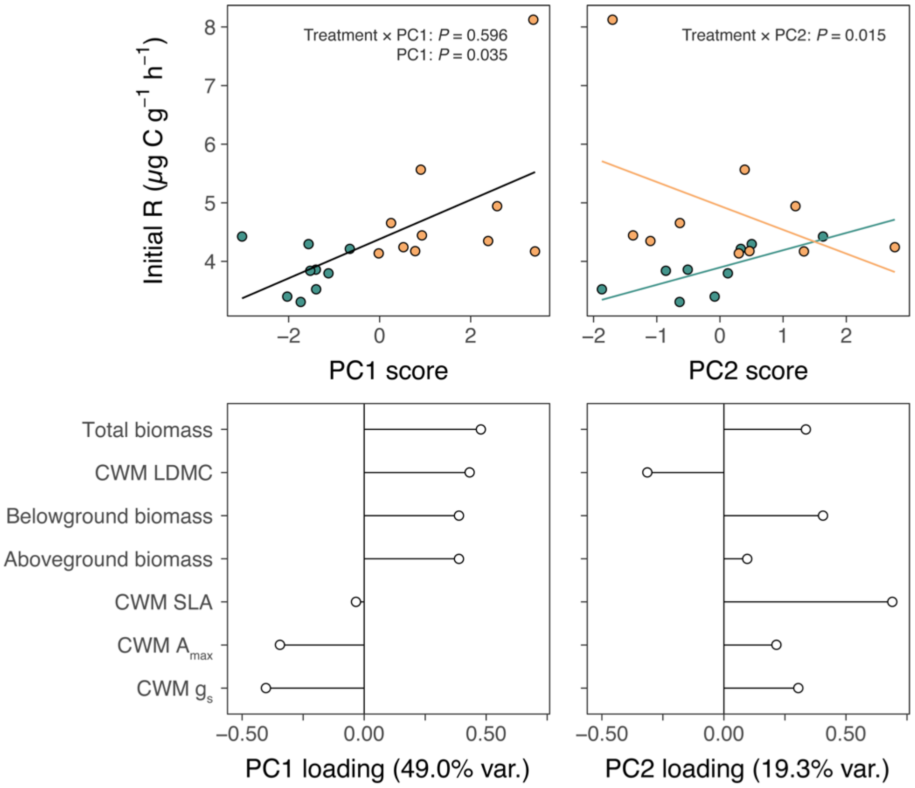

**Fig. S4 | Alpine and lowland plant effects on alpine soil microbial respiration.** (a,b) Relationships between plant physiology, expressed as (a) PC1 and (b) PC2 scores from PCAs of pot-level measurements, and rates of soil microbial respiration during the first nine days of incubation (initial R; μg C $g^{-1}$- $h^{-1}$) in alpine (green) versus lowland (yellow) treatments of the glasshouse experiment. Fit lines describe relationships as determined by a single linear mixed effects model including treatment, PC1, PC2 and treatment × PC# interactions (N = 20; Main Text), with different colours in (b) indicating a significant treatment × PC2 interaction. (c,d) Loadings of plant physiological measurements on (c) PC1 and (d) PC2 axes. Total, belowground and aboveground biomass are sums of all plants per pot and specific leaf area (SLA), maximum photosynthetic capacity ($A_{max}$) and stomatal conductance ($g_s$) are pot-wise community weighted mean (CWM) values.

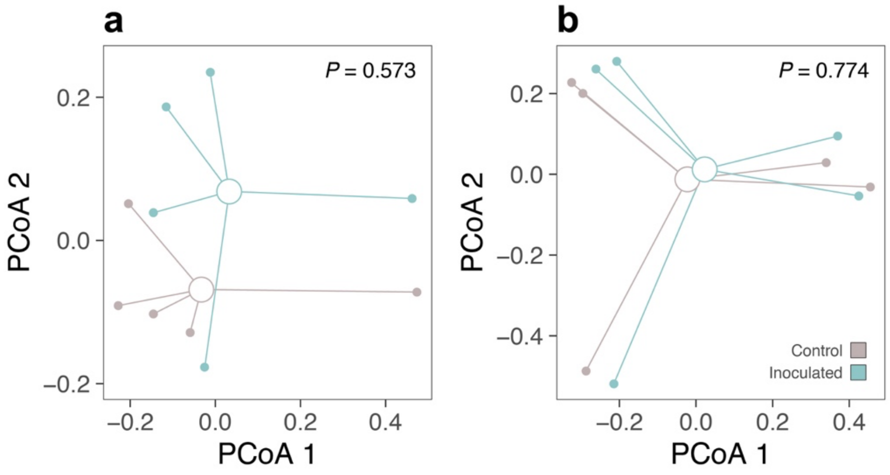

**Fig. S5 | Soil inoculation effects on alpine soil community composition.** Effects of low elevation soil biota inoculation on (a) bacterial and (b) fungal community composition in alpine plots (N = 10 in both cases). Axes show the first two components of principal coordinates analyses (PCoA; Bray Curtis distance) performed on the relative abundances of bacterial and fungal operational taxonomic units, respectively. *P*-values are from PERMANOVAs testing for inoculation effects on the same distance matrices.

**Table S1. Linear mixed effects model outputs for effects of field treatment (control, warming, warming plus lowland plants), region and their interaction on soil variables.**

| Response | Treatment | | Region | | Treatment × Region | |
|---|---|---|---|---|---|---|
| | LR | *P* | LR | *P* | LR | *P* |
| Soil carbon content | 23.16 | < 0.0001 | 34.53 | < 0.0001 | 4.64 | 0.0984 |
| Ecosystem respiration | 49.61 | < 0.0001 | 13.08 | 0.0045 | 12.98 | 0.0015 |
| Microbial biomass C | 33.52 | < 0.0001 | 13.64 | 0.0034 | 5.35 | 0.0688 |
| Microbial growth (per gram soil) | 9.33 | 0.0094 | - | - | - | - |
| Microbial respiration (per gram soil) | 1.20 | 0.5500 | - | - | - | - |
| Microbial growth (biomass-specific) | 8.32 | 0.0156 | - | - | - | - |
| Microbial Respiration (biomass-specific) | 6.54 | 0.0381 | - | - | - | - |
| Microbial carbon use efficiency | 15.68 | 0.0004 | - | - | - | - |

**Table S2. Linear mixed effects model outputs for treatment (alpine plants, lowland plants) effects on soil variables in the glasshouse experiment.**

| Response variable | LR | *P* |
|---|---|---|
| Soil pore water absorbance ($a_{350}$) | 24.84 | < 0.0001 |
| Soil pore water total fluorescence ($F_{tot}$) | 27.72 | < 0.0001 |
| Soil pore water fluorescence index (FI) | 9.55 | 0.0020 |
| Microbial biomass carbon | 0.16 | 0.6932 |
| Fast-decaying soil carbon pool size | 3.99 | 0.0458 |
| Fast-decaying soil carbon pool decay rate | 0.23 | 0.6306 |
| Soil DOM C1 (protein-like) | 21.39 | < 0.0001 |
| Soil DOM C2 (protein-like) | 10.42 | 0.0012 |
| Soil DOM C3 (humic-like) | 15.65 | 0.0001 |
| Soil DOM C4 (humic-like) | 9.60 | 0.0019 |
| Soil DOM C5 (humic-like) | 17.39 | < 0.0001 |
| Soil DOM C6 (fulvic acid-like) | 16.50 | < 0.0001 |

**Table S3. Statistical test outputs for lowland species identity effects on soil carbon loss.**

| **Explanatory variable** | **Linear model test statistics** | | |
|---|---|---|---|
| | F | d.f. | *P* |
| Western Alps experiment | | | |
| *Achillea millefolium* | 0.17 | 1,4 | 0.7007 |
| *Bellis perennis* | 0.40 | 1,4 | 0.5634 |
| *Bromus erectus*[*] | - | - | - |
| *Dactylis glomerata*[*] | - | - | - |
| *Medicago lupulina*[*] | - | - | - |
| *Plantago media* | 1.19 | 1,4 | 0.3364 |
| *Salvia pratensis* | 0.10 | 1,4 | 0.7684 |
| Central Alps experiment | | | |
| *Brachypodium pinnatum* | 3.70 | 1,8 | 0.3053 |
| *Carex flacca* | 28.24 | 1,8 | 0.1184 |
| *Carum carvi* | 5.94 | 1,8 | 0.2479 |
| *Dactylis glomerata*[*] | - | - | - |
| *Hypericum perforatum* | 10.11 | 1,8 | 0.1939 |
| *Plantago lanceolata*[*] | - | - | - |
| *Primula veris* | 6.41 | 1,8 | 0.2395 |
| *Ranunculus bulbosus* | 40.38 | 1,8 | 0.0994 |
| *Salvia pratensis* | 0.08 | 1,8 | 0.8281 |
| *Silene vulgaris* | 36.12 | 1,8 | 0.1050 |
| *Trifolium montanum*[*] | - | - | - |
| *Viola hirta*[*] | - | - | - |

[*] *NB: species not tested, present in < 3 plots*

**Table S4. DOM components identified through PARAFAC modelling.**

| Name | Excitation λ (nm) | Emission Λ (nm) | General description[†] | Peak name[*] |
|---|---|---|---|---|
| C1 | 275 | 332 | Protein-like (tryptophan) | T |
| C2 | 270 | 310 | Protein-like (tyrosine) | B |
| C3 | 340 | 434 | Humic-like | A/C |
| C4 | 305 | 424 | Microbial humic-like | M |
| C5 | 375 | 452 | Humic-like | C |
| C6 | < 250 (400) | 506 | Soil fluvic acid-like | C+ |

[*] Ref (*10*); [†] Ref (*9*)
